# Adaptation of anammox bacteria to low temperature via gradual acclimation and cold shocks: distinctions in protein expression, membrane composition and activities

**DOI:** 10.1101/2021.08.29.458117

**Authors:** V. Kouba, D. Vejmelkova, E. Zwolsman, K. Hurkova, K. Navratilova, M. Laureni, P. Vodickova, T. Podzimek, J. Hajslova, M. Pabst, M.C.M. van Loosdrecht, J. Bartacek, P. Lipovova, D.G. Weissbrodt

## Abstract

Anammox bacteria enable an efficient removal of nitrogen from sewage in processes involving partial nitritation and anammox (PN/A) or nitrification, partial denitrification, and anammox (N-PdN/A). In mild climates, anammox bacteria must be adapted to ≤15 °C, typically by gradual temperature decrease; however, this takes months or years. To reduce the time necessary for the adaptation, an unconventional method of ‘cold shocks’ is promising, involving hours-long exposure of anammox biomass to extremely low temperatures. We compared the efficacies of gradual temperature decrease and cold shocks to increase the metabolic activity of anammox (fed batch reactor, planktonic “*Ca*. Kuenenia”). We assessed the cold shock mechanism on the level of protein expression (quantitative shot-gun proteomics, LC-HRMS/MS) and structure of membrane lipids (UPLC-HRMS/MS). The shocked culture was more active (0.66±0.06 vs 0.48±0.06 kg-N/kg-VSS/d) and maintained the relative content of N-respiration proteins at levels consistent levels with the initial state, whereas the content of these proteins decreased in gradually acclimated culture. Cold shocks also induced a more efficient up-regulation of cold shock proteins (e.g. CspB, TypA, ppiD). Ladderane lipids characteristic for anammox evolved to a similar end-point in both cultures which confirms their role in anammox bacteria adaptation to cold and indicates a three-pronged adaptation mechanism involving ladderane lipids (ladderane alkyl length, introduction of shorter non-ladderane alkyls, polar headgroup). Overall, we show the outstanding potential of cold shocks for low-temperature adaptation of anammox bacteria and provide yet unreported detailed mechanisms of anammox adaptation to low temperatures.

**Highlights:**

- Anammox bacteria were adapted to low T by gradual acclimation and cold shocks
- The shocked culture was more active (0.66±0.06 vs 0.48±0.06 kg-N/kg-VSS/d)
- N-respiration proteins content decreased in gradually acclimated bacteria
- Several cold shock proteins were upregulated more efficiently by cold shocks
- At ↓T, anammox adjusted ladderane membrane lipid composition in three aspects

**Graphical abstract:** 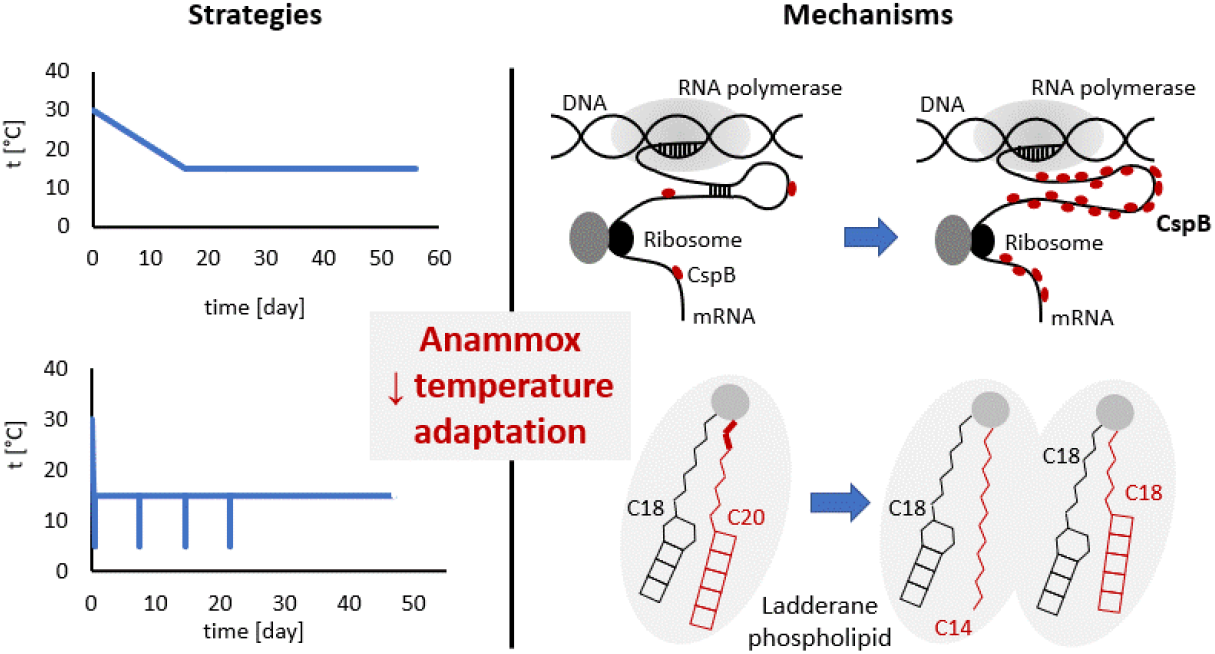

## 1 Introduction

Anammox bacteria can cost-effectively remove nitrogen from the mainstream of WWTP via processes such as partial nitritation and anammox (PN/A) as well as nitrification, partial denitrification and anammox (N-PdN/A). However, the mainstream of WWTP is much colder (10-20 °C) than most current anammox reactors used to treat mesophilic centrates (30-37 °C), making the adaptation of anammox bacteria to low temperatures a crucial issue(Cao et al., 2017; Kouba et al.).

Although some studies have reported that anammox bacteria deteriorated in activity at ≤20 °C (Hoekstra et al., 2018), others have shown that anammox adapted and culture activity increased (Lv et al., 2020; Wang et al., 2018). This activity can be further elevated by the enrichment of cold-adapted species (Hendrickx et al., 2014; Park et al., 2017), by the gradual acclimation of mesophilic cultures to a psychrophilic regime (De Cocker et al., 2018) or by short-term exposure to extremely low temperature called the cold shock (Kouba et al., 2018).

Different cold-adaptation strategies have been applied to enhance anammox activities at low temperatures. First, the enrichment of cold-adapted anammox has been initiated with activated sludge and after 700 days of cultivation at 10 °C, relevant activities of 30-44 g-N/kg-VSS/d have been achieved (Hendrickx et al., 2014). Second, a gradual acclimation strategy has involved a stepwise reduction of cultivation temperature from 30 to 20, 15, 12.5 and finally 10 °C over the period of 349 days, after which an impressive activity of 91.8 g-Namon/kg-VSS/d has been achieved (De Cocker et al., 2018). Third, cold shock strategy has included a rapid cooling of anammox biomass (“*Candidatus* Brocadia”) from 24 °C to 5 °C, then 8 hours of exposure to 5 °C, after which the temperature has been raised back to 24 °C. Three such shocks applied over the period of 45 days have incrementally increased the anammox activity at 10 °C to a relevant 54 g-N/kg-VSS/d, compared to an order of magnitude less active control (Kouba et al., 2018). Most recently, a single cold shock (5 °C, 8 h) doubled anammox (“*Ca*. Brocadia” and “*Ca*. Scalindua”) activity at 15 °C for at least 40 days, indicating a highly promising long-term adaptive effect (Kouba et al., accepted). However, these and actually most other studies of low-temperature anammox restrict their focus to cultures activities and dominant anammox populations. The absence of comparison raises questions about their relative efficiency and precise mechanisms of action.

Some researchers have attempted to explain the differences between the aforementioned cold adaptation strategies and their specific mechanisms in physiological terms. For example, (Rattray et al., 2010), reporting on the membrane composition of “*Ca*. Scalindua”, showed that the length of a single ladderane lipid alkyl (C18/C20 [5]-ladderane ester) increased as the cultivation temperature increased from 10 to 20 °C. Recently, membrane composition was correlated to anammox temperature coefficients; specifically, higher relative contents of short C18 [3]-ladderane alkyls and large phosphatidylcholine headgroups were detected in cultures more active at 15-30 °C and 10-15 °C, respectively (Kouba et al.). This shows that other lipid structures are also a key part of the mechanism involved in anammox adaptation to low temperatures. Furthermore, only two metaproteomic studies assessing anammox low-temperature adaptation are available. In the first one, changes in the protein content indicated a distinct inhibition as the temperature decreased from 35 to 15 °C (Lin et al., 2018). Of the three anammox genera detected in the latter study, “*Ca*. Jettenia” displayed the most changes in protein expression, suggesting that certain anammox genera may have advantages at low temperatures. In the second study, a reducing the temperature of anammox culture from 35 to 25 °C appeared to stimulate the upregulation of cold shock proteins, potentially giving “*Candidatus* B. fulgida” an advantage over “*Ca*. B. sinica” and “*Ca*. Jettenia caeni” (Huo et al., 2020); however, these proteins were unspecified. Despite these advances, a lot remains to be known about the various mechanisms of cold adaptation.

To further such knowledge, we tested the adaptation of a mesophilic culture of “*Ca*. Kuenenia stuttgartiensis” to main-stream temperatures by the application of either a gradual cold acclimation strategy (1 °C per day) or cold shocks (duration 1 h, temperature 5 °C). The strategies were compared in terms of anammox activity, protein expression via large-scale metaproteomics, and membrane composition including ladderane phospholipids. Specifically, we aimed to understand how anammox bacteria adapt their metabolism to cold temperature and what acclimation regime can help maintain substantial anammox activities at low temperatures.

## 2 Materials and Methods

### 2.1 Inoculum

Planktonic biomass enriched in “*Candidatus* Kuenenia stuttgartiensis” was adopted as inoculum. According to a 16S rRNA gene amplicon sequencing, the enrichment degree was 76%. Within family Brocadiaceae “*Ca*. Kuenenia” covered 97 % and the rest was represented by “*Ca*. Brocadia (2 %), and “*Ca*. Scalindua (1 %) (Kouba et al., nonpublished data). The inoculum was obtained from a membrane bioreactor (MBR) maintained at 30 °C and a pH of 7, under which the doubling time was 3 days. The details of seeding reactor operation are described in (Hoekstra et al., 2018).

### 2.2 Bioreactor operation

Three fed batch reactors (FBRs) (effective volume 1 L, Fig. S 1) were seeded by inoculum from an anammox continuous-flow membrane bioreactor (MBR). The FBRs were kept anoxic by flushing the liquid phase with the mixture of N2/CO2 (95/5%, CO2 to maintain carbonate equilibrium in reactor liquid phase and provide CO2 for anabolism). Table 1 shows the description of all FBRs. After an initial test of anammox activity at 30 °C was performed, the temperature in control FBR was reduced gradually from 30 to 15 °C at a rate of 1 °C per day, after which it remained at 15 °C for 34 days. In the second FBR, after the initial activity test at 30 °C, the reactor was cooled down from 30 to 5 °C in 10 minutes and after 60 min at 5 °C, it was warmed up to 15 °C at the same speed as it was cooled down (1.5 °C/min). Afterwards it was shocked to 5 °C every 3 days (8 shocks in total). The third FBR was operated identically as the second one, except that the shocks were performed every 7 days (4 shocks in total). In the 2nd and 3rd reactor, the period with shocks was followed by a consistent operation at 15 °C.

**Table 1:** Description of operated fed batch reactors.

|  |  |
| --- | --- |
| Gradual decrease<br>(Control) | Temperature decreased from 30 to 15 °C over 15 days (Control), 1 °C per day |
| Shocked every 3 days | 1st cold shock:<br>17 min: 30 °C → 5 °C; 60 min: 5 °C; 7 min: 5 °C → 15 °C<br>→ operation at 15 °C until next shock<br>2nd-8th cold shock:<br>17 min: 15 °C → 5 °C; 60 min: 5 °C; 7 min: 5 °C → 15 °C<br>→ operation at 15 °C until next shock |
| Shocked every 7 days | 1st cold shock:<br>17 min: 30 °C → 5 °C; 60 min: 5 °C; 7 min: 5 °C → 15 °C<br>→ operation at 15 °C until next shock<br>2nd-4th cold shock:<br>17 min: 15 °C → 5 °C; 60 min: 5 °C; 7 min: 5 °C → 15 °C<br>→ operation at 15 °C until next shock |

To perform the cold-shocks, a water-ice-NaCl cooling bath with the ratio of 10:10:1 kg, respectively was used. This ratio resulted in a fluid temperature of −4 °C which ensured fast cooling down as well as stopping at 5 °C to prevent reaching lower temperature. A polystyrene box was used as an insulating container for the cooling bath.

The FBRs were fed by ammonium (230 g NH4Cl/L, 60 mgN-NH_4_^+^/mL) and nitrite solutions (296 g NaNO2/L, 60 mgN-NO2-/mL). Other constituents of the media are described in Tables S 1-2. The FBRs were operated with excess ammonium (100-200 mgN-NH4+/L), and nitrite was spiked to 30 mg-N-NO2-/L as often as it was consumed to prevent long starvation.

Anammox activity was determined by sampling throughout the FBR cycle and analyzing the concentration of ammonium and nitrite nitrogen spectrophotometrically on the Gallery™ Discrete Analyzer (Thermo Fisher Scientific) according to the standard methods (Apha, 2005). The concentrations of ammonium and nitrite over time were fitted with linear regressions, whose slopes were determined as volumetric removal rates. The biomass concentration was determined by way of optical density (OD) at 600 nm and 660 nm using 1 mL of biomass mixture, recalculated to volatile suspended solids. Specific anammox activity was expressed as the sum of ammonium and nitrite removal rate per mass of volatile suspended solids and time. Activation energies were calculated according to Arrhenius (equations 1 and 2), where k is the ratio of anammox activities at the lower (numerator) and higher (denominator) compared temperatures, *ln* is the natural logarithm, *A* is a constant pre-exponential factor, *Ea* is the activation energy (J.mol^-1^), *R* is the ideal gas constant (J.mol.^-1^.K^-1^) and *T* is the thermodynamic temperature (K).

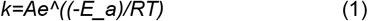

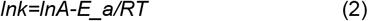

### 2.3 Label free quantification (LFQ) by shot-gun proteomics

A volume of 2 mL of biomass was centrifuged at 14 000 x g at 4 °C for 3 min. The supernatant was discarded and the pellet was stored at – 20 °C until further analysis.

#### 2.3.1 Protein extraction

All samples were processed in duplicates. 350 μL of TEAB Resuspension buffer and 350 μL of B-PER buffer was added to the pellet. After resuspension 140 mg of glass beads (Sigma-Aldrich, 150-212 μm, 70-100 U.S. sieve) were added and the mixture was subjected to bead beating: 30 s five times with 30 s on ice in between. Then three freeze/thaw cycles were applied (liquid nitrogen/80 °C for 3 min). The sample was centrifuged at 14 000 x g, 4 °C, 10 min. 0.5 mL of the supernatant was mixed with 125 μL 100% (w/v) TCA, vortexed and incubated on ice for 10 min.

The tube was centrifuged at 14 000 x g at 4 °C for 5 min. The supernatant was removed and the pellet was washed two times with cold acetone (200 μL of cold acetone, centrifugation 14 000 x g at 4 °C for 5 min). The supernatant was removed and 300 μL of 6M Urea in 200 mM ammonium bicarbonate (ABC). After homogenization 90 μL of 10mM DTT was added and the tube was incubated at 37 °C for 60 min. Then 90 μL of 20mM IAM in ABC was added and the tube was incubated at RT in the dark for 30 min. The mixture was vortexed and spinned down. 160 μL of supernatant was mixed with 240 μL 200mM ABC. After vortexing 5 μL of trypsin was added, vortexed again and the mixture was digested overnight at 37 °C, 300 rpm in the dark.

#### 2.3.2 Large-scale shot-gun metaproteomics

An aliquot corresponding to approx. 250ng protein digest was analysed from each duplicate preparation using a one dimensional shot-gun proteomics approach (Köcher et al., 2012). Briefly, the samples were analysed using a nano-liquid-chromatography system consisting of an EASY nano LC 1200, equipped with an Acclaim PepMap RSLC RP C18 separation column (50 μm x 150 mm, 2μm), and an QE plus Orbitrap mass spectrometer (Thermo). The flow rate was maintained at 350 nL/min over a linear gradient from 5% to 25% solvent B over 88 minutes, and finally to 50% B over 25 minutes, followed by back equilibration to starting conditions. Data were acquired from 2 to 119 min. Solvent A was H_2_O containing 0.1% formic acid, and solvent B consisted of 80% acetonitrile in H_2_O and 0.1% formic acid. The Orbitrap was operated in data-dependent acquisition mode acquiring peptide signals form 385-1250 m/z at 70K resolution with a max IT of 100ms and an AGC target of 3e6. The top 10 signals were isolated at a window of 2.0 m/z and fragmented using a NCE of 28. Fragments were acquired at 17K resolution with a max IT of 100ms and an AGC target of 5e4.

#### 2.3.3 Database search, label free quantification and visualisation

Data were analysed against the proteome database from “*Candidatus* Kuenenia stuttgartiensis (UniprotKB, Kuenenia stuttgartiensis, TaxID 174633) using PEAKS Studio 10.0 (Bioinformatics Solutions Inc)2 allowing for 20 ppm parent ion and 0.02 m/z fragment ion mass error, 2 missed cleavages, carbamidomethylation as fixed and methionine oxidation and N/Q deamidation as variable modifications. Peptide spectrum matches were filtered against 1% false discovery rate (FDR) and protein identifications with ≥ 2 unique peptides were accepted. Changes between individual conditions were further evaluated using the label free quantification (LFQ) option provided by the PEAKS Q software tool (Bioinformatics Solutions Inc). A pairwise comparison of the above described conditions was performed on identified peptide spectra filtered against 1% FDR, a mass error equal or less to 12.5 ppm and a max. RT shift between runs of 1.5 minutes. Peptides with variable modifications were excluded.

The significance method was set to ANOVA with a significance level threshold of ≥13, 1.5 fold change and 2 unique peptides per protein. Additional peptide filters for visualisation were set for spectral quality to equal or greater 2, average intensity to equal or greater 1E4, charge states were restricted to between 1 to 6, and the limit of confident samples and peptide ID counts were set to 0. Data were further visualised in hierarchical clustered protein profile heatmaps (Välikangas et al., 2017)

### 2.4 Ladderane analysis

We used UPLC-HRMS/MS which is exceptionally sensitive and provided insight into the number of carbon atoms of these lipids. Per (Rattray et al., 2008), the polar headgroup ionization differs substantially, so the results can be characterized only qualitatively.

#### 2.4.1 Reagents and chemicals

Deionized water was obtained from a Milli-Q^®^ Integral system supplied by Merck (Darmstadt, Germany). HPLC-grade methanol, isopropyl alcohol, formic acid and ammonium formate (purity ≥ 99 %) were purchased from Sigma-Aldrich (St. Luis, MO, USA).

#### 2.4.2 Sample preparation

To extract ladderane phospholipids, a mixture of MeOH:DCM:10mM ammonium acetate (2:1:0.8, v/v/v) was chosen according to Lanekoff & Karlsson (2010). Lyophilized anammox cultures were weighted (0.2 g) into a plastic cuvette and automatically shaken for 2 min with 2 mL of extraction solvent. The suspensions were sonicated for 10 min, centrifuged (5 min, 10000 rpm, 5 °C). Finally, 1 mL of supernatant was transferred into the vial before further analysis by ultra-high performance liquid chromatography coupled to high-resolution tandem mass spectrometry (U-HPLC-HRMS/MS).

#### 2.4.3 Ultra-high performance liquid chromatography coupled to high-resolution mass spectrometry (U-HPLC-HRMS)

The Dionex UltiMate 3000 RS U-HPLC system (Thermo Fisher Scientific, Waltham, USA) coupled to quadrupole-time-of-flight SCIEX TripleTOF^®^ 6600 mass spectrometer (SCIEX, Concord, ON, Canada) was used to analyse ladderane phospholipids. Chromatographic separation of extracts was carried out using U-HPLC system, which was equipped with Acquity UPLC BEH C18 column, 100Å, 100 mm × 2.1 mm; 1.7 μm particles (Waters, Milford, MA, USA). The mobile phase consisted of (A) 5 mM ammonium formate in Milli-Q water:methanol with 0.1% formic acid (95:5 v/v) and (B) 5 mM ammonium formate in isopropyl alcohol:methanol: Milli-Q water with 0.1% formic acid (65:30:5, v/v/v).

The following elution gradient was used in positive ionization mode: 0.0 min (90% A; 0.40 mL min^-1^), 2.0 min (50% A; 0.40 mL min^-1^), 7.0 min (20% A; 0.40 mL min^-1^), 13.0 min (0% A; 0.40 mL min^-1^), 20.0 min (0% A; 0.40 mL min^-1^), 20.1 min (95% A; 0.40 mL min^-1^), 22.0 min (90% A; 0.40 mL min^-1^).

The sample injection volume was set at 2 μL, the column temperature was kept constant at 60 °C and autosampler temperature was permanently set at 5 °C. A quadrupole-time-of-flight TripleTOF^®^ 6600 mass spectrometer (SCIEX, Concord, ON, Canada) was used. The ion source Duo Spray™ with separated ESI ion source and atmospheric-pressure chemical ionization (APCI) was employed. In the positive ESI mode, the source parameters were set to: nebulizing gas pressure: 55 psi; drying gas pressure: 55 psi; curtain gas 35 psi, capillary voltage: +4500 V, temperature: 500 °C and declustering potential: 80 V.

The other aspects of the methodology were consistent with (Hurkova et al., 2019), except for the confirmation of compound identification, which used accurate mass, isotopic pattern and MS/MS characteristic fragments.

## 3 Results and Discussion

### 3.1 Cultures exposed to cold shocks achieved higher anammox activity than a gradually acclimated culture

Initial batch activity tests at 30 °C showed that the metabolic activities of the three anammox cultures (two shocked, one gradually acclimated) compared favorably (2.60-2.88 kg-N/kg-VSS/d, Fig. 1). Subsequently, the shocked cultures were exposed to 5 °C for 1 h every 3 or 7 days (15 °C between each shock; shocks concluded on days 26 and 22, respectively) while the temperature of the gradually acclimated culture was reduced by 1 °C per day. During the cold shocks anammox activity was not detected. Between shocks (15 °C), the cultures that were shocked every 3 or 7 days had an average activity of 0.81±0.12 and 0.73±0.08 kg-N/kg-VSS/d (average±standard deviation), respectively. The activity of the gradually acclimated culture at 30, 25, 20 and 15 °C was 2.8, 2.0, 1.2 and 0.45±0.06 kg-N/kg-VSS/d (average±standard deviation), respectively.

**Fig. 1:**
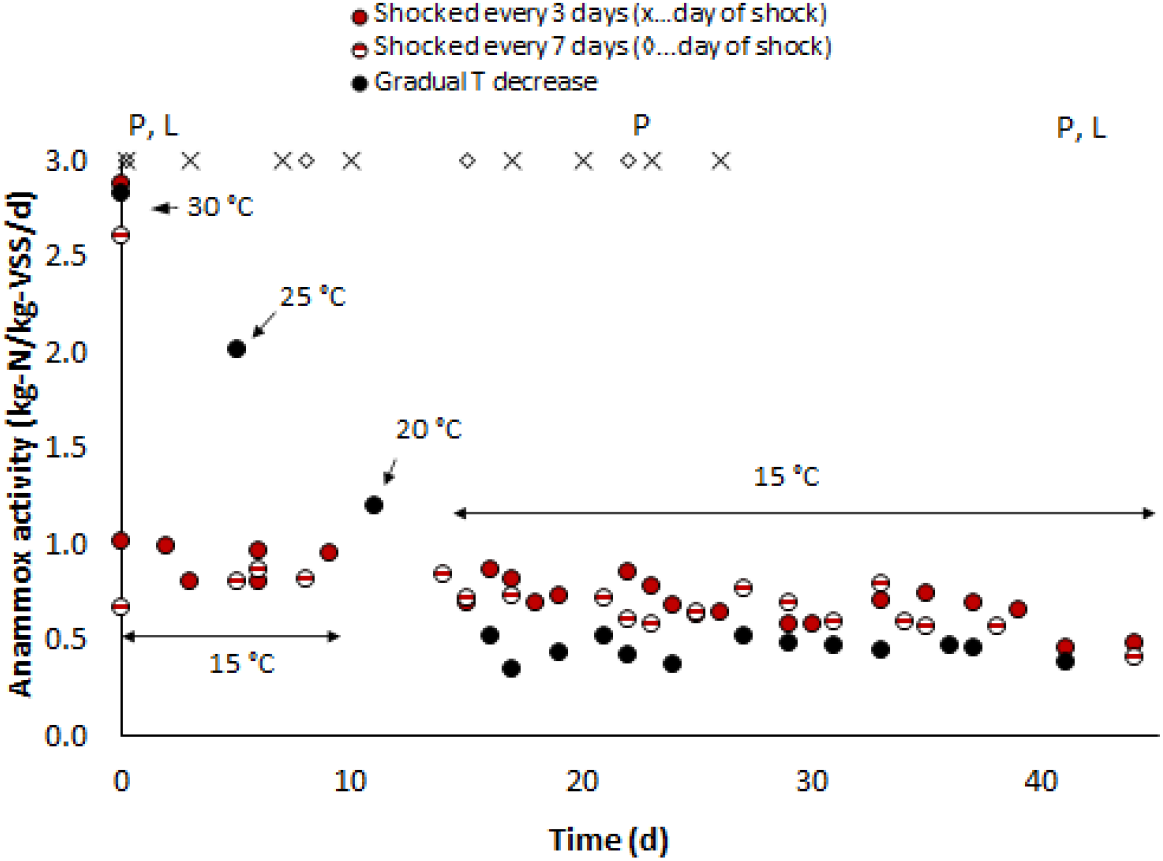
Comparison of strategies for the adaptation of planktonic anammox bacteria (“Ca. Kuenenia”) from mesophilic (30 °C) to psychrophilic (15 °C) temperatures. At 15°C, anammox biomass exposed to cold shocks (5 °C, 1 h) were more active compared to biomass acclimatized by gradual temperature decrease (1 °C per day). P and L timepoints indicate sampling for protein and lipid analyses.

After the termination of the adaptation regimes (shocks and gradual acclimation), their long-term effect on the activities of the cultures was assessed at 15 °C. Interestingly, the cultures shocked every 3 or 7 days had similar activities of 0.66±0.06 and 0.65±0.10 kg-N/kg-VSS/d, respectively, whereas the gradually acclimated culture had a significantly lower activity of 0.48±0.02 kg-N/kg-VSS/d. The shocked cultures were by 36% more active than the gradually acclimated culture. In this post-adaptation phase, the temperature coefficients for 15-30 °C in the shocked cultures were similar (66 kJ/mol) and lower than the corresponding coefficient for the gradually acclimated culture (86 kJ/mol). The activities of all cultures converged on day 40; this was 14 and 18 days (cultures shocked every 3 and 7 days, respectively) after the last shock was concluded for each culture; specifically, the activities of the shocked cultures decreased to that of the level of the acclimated culture. Overall, this shows that the cold shocks induced the more efficient adaptation of anammox to 15 °C compared to the gradual acclimation, if only for 14-18 days.

### 3.2 Protein expression explains the lower activity of acclimated anammox and underlying mechanisms of cold adaptation

Protein expression in two of the cultures (shocked every 7 days, gradually acclimated) was determined using large-scale shotgun metaproteomics: (i) in the initial sample at 30 °C, (ii) after the conclusion of the adaptation regimes, during which the shocked culture was more active than the acclimated culture (15 °C), and (iii) after the activities of the cultures converged.

#### 3.2.1 Proteins involved in the nitrogen metabolism of anammox bacteria

The lower activity of the gradually acclimated anammox was linked to the relative content of the proteins responsible for nitrogen conversion, such as nitrate oxidoreductase (NXR), nitrite reductase (Nir), hydrazine synthase (HZS), hydroxylamine oxidoreductase (HAO) and hydrazine dehydrogenase (HDH). In the gradually acclimated culture, the content of these proteins decreased compared to initial baseline (Table 2). Most reduced were α and β subunits of Nir, α and γ subunits of HZS, and HDH. Conversely, in the shocked culture, the content of these proteins did not change as significantly. This suggests that the lower activity of the gradually acclimated culture was due to the reduced content of several metabolic proteins.

**Table 2:**
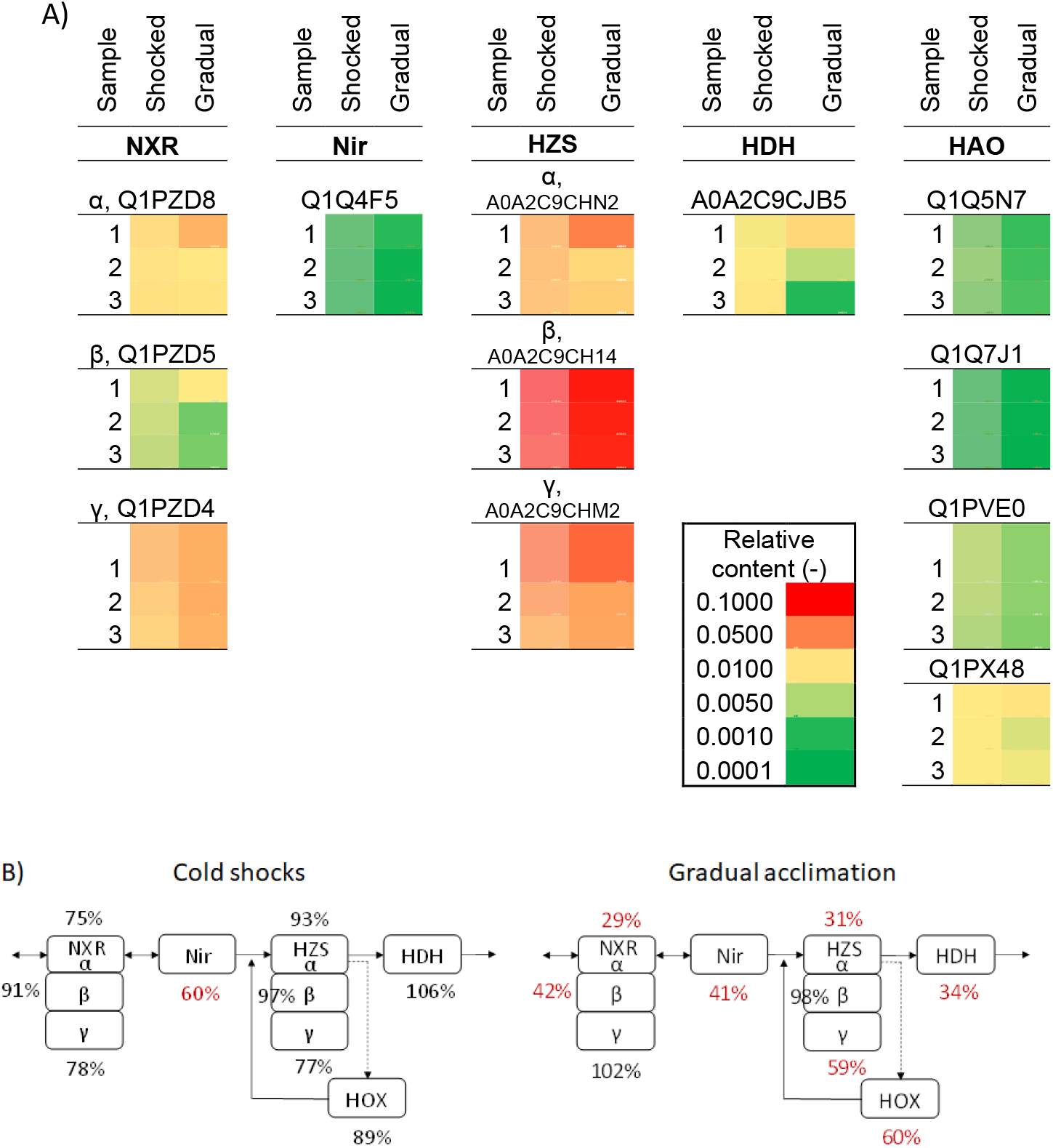
Relative content of proteins of anammox nitrogen respiration in biomasses exposed to cold shocks and gradual acclimation. Sample 1 - day 0, sample 2 - divergent activities of shocked and gradually acclimated cultures, sample 3 - end of experiment and cultures activities converged (schematic A). The schematic B shows % of metabolic protein content in sample 2 compared to initial state (day 0), indicating that the content of these proteins decreased much more significantly in the gradually acclimated biomass.

The content of metabolic proteins was also linked to the decreased activity of the shocked culture 21 days post-shock. In this period, several anammox metabolic proteins (HZS, NXR and HAO) decreased in abundance in the shocked culture while remaining the same in the gradually acclimated one. Most pronounced decrease was noted for Nir that decreased (relative protein content at the end of experiment compared to immediately post-shock, samples 2 and 3, Table 2) compared to unchanged content in the gradually acclimated culture.

#### 3.2.2 Role of cold shock proteins in anammox low-temperature adaptation

To explain the mechanism of cold adaptation, label-free quantification was used to identify the most up- or down-regulated proteins. We attributed the most relevance to those matching the expression profile described by (Horn et al., 2007), proteins that were both upregulated by cold shocks and then down-regulated after the activities of the cultures converged. Only 9 proteins matched this description (Table 3), including the cold shock protein B (CspB). Compared to the initial state, cold shocks temporarily upregulated CspB (shocked 5-fold; gradually acclimated 12-fold) to a slightly higher relative content compared to gradual acclimation (9.5E-03 vs. 8.9E-03). The eight other suspected cold shock proteins were less abundant than CspB and either more efficiently upregulated or more abundant in the shocked culture than in the gradually acclimated one. Furthermore, the proteomics experiment suggested that 21 other proteins (Table 4) were more efficiently upregulated by cold shocks, bringing the total number of proteins potentially involved in the low-temperature adaptation of anammox by cold shocks to 30.

**Table 3:** Comparison of proteins expressions in anammox biomasses adapted to low temperature (15°C) via cold shocks and via gradual acclimation. Proteins upregulated immediately after shocks and downregulated after period without the shocks. Shocks induced more efficient upregulation than acclimation. Sample 1 - day 0, sample 2 - divergent activities of shocked and gradually acclimated cultures, sample 3 - end of experiment and cultures activities converged.

| Description, gene tag, accession number | Sample | Relative content |  | Fold change compared to initial state |  |
| --- | --- | --- | --- | --- | --- |
|  |  | Shocked | Gradual | Shocked | Gradual |
| Strongly similar to cold shock protein CspB, KSMBR1_1099, Q1Q191 (or Q1PVN1) | 1 |  |  |  |  |
|  | 2 |  |  | 5.2 | 12 |
|  | 3 |  |  |  |  |
| Strongly similar to GTP-binding protein TypA, KSMBR1_0377, Q1Q4Q3 | 1 |  | n.d. |  |  |
|  | 2 |  |  | 6.3 | 53 |
|  | 3 |  |  |  |  |
| Periplasmic folding chaperone, ppid, Q1Q206 | 1 |  |  |  |  |
|  | 2 |  |  | 3.2 | 1.4 |
|  | 3 |  |  |  |  |
| Uncharacterized protein, kustd1530, Q1PYW7 | 1 |  |  |  |  |
|  | 2 |  |  | 3.0 | 2.3 |
|  | 3 |  |  |  |  |
| Universal stress protein, UspA, Q1PWE5 | 1 |  |  |  |  |
|  | 2 |  |  | 2.6 | 1.5 |
|  | 3 |  |  |  |  |
| Similar to PQQ biosynthesis protein C, pqqC, Q1PYI1 | 1 |  |  |  |  |
|  | 2 |  |  | 2.5 | 2.1 |
|  | 3 |  |  |  |  |
| Similar to heme d1 synthesis protein, NirD/nirG, Q1PXI2 | 1 |  |  |  |  |
|  | 2 |  |  | 2.5 | 1.9 |
|  | 3 |  |  |  |  |
| Methionine aminopeptidase, map, Q1Q155 | 1 |  | n.d. |  |  |
|  | 2 |  | n.d. | 2.5 |  |
| 30S ribosomal protein S18, rpsR, Q1Q7I5 | 1 |  |  |  |  |
|  | 2 |  |  | 1.2 | 1.2 |
|  | 3 |  |  |  |  |

**Table 4 A:** Comparison of anammox proteins expressions in anammox biomasses adapted to low temperature (15°C) via cold shocks and via gradual acclimation. Anammox proteins upregulated more efficiently by cold shocks, excluding proteins already mentioned in Table 3. Sample 1 - day 0, sample 2 - distinct activities, sample 3 - end of experiment. Note: Red triangle in upper right corner…protein not detected, the value was set to detection limit 1E-6.

| Description, gene tag, accession number | Sample | Relative content |  | Fold change compared to initial state |  |
| --- | --- | --- | --- | --- | --- |
|  |  | Shocked | Gradual | Shocked | Gradual |
| Hemerythrin HHE cation binding domain protein, KSMBR1_0022, A0A2C9C9T9 | 1 |  |  |  |  |
|  | 2 |  |  | 52 | 8 |
|  | 3 |  |  |  |  |
| Selenate reductase assembly chaperone protein, serD, A0A2C9CE34 | 1 |  |  |  |  |
|  | 2 |  |  | 16 | 5 |
|  | 3 |  |  |  |  |
| Similar to thiS (Thiamine biosynthesis), thiS, Q1PW59 | 1 |  |  |  |  |
|  | 2 |  |  | 5.9 | 1.9 |
|  | 3 |  |  |  |  |
| Uncharacterized protein, KSMBR1_3335, A0A2C9CJH3 | 1 |  |  |  |  |
|  | 2 |  |  | 5.2 | 1.8 |
|  | 3 |  |  |  |  |
| Uncharacterized protein, kuste2355, Q1Q698 | 1 |  |  |  |  |
|  | 2 |  |  | 5.2 | 1.8 |
|  | 3 |  |  |  |  |
| Proline--tRNA ligase, proS, Q1Q7M8 | 1 |  |  |  |  |
|  | 2 |  |  | 4.7 | 0.9 |
|  | 3 |  |  |  |  |
| Proline--tRNA ligase, proS, A0A2C9CIY0 | 1 |  |  |  |  |
|  | 2 |  |  | 4.7 | 0.9 |
|  | 3 |  |  |  |  |
| UDP-glucose 6-dehydrogenase, KSMBR1_0073, Q1Q302 | 1 |  |  |  |  |
|  | 2 |  |  | 4.3 | 1.2 |
|  | 3 |  |  |  |  |
| Anti-sigma factor RsrA, rsrA, Q1Q6G9 | 1 |  |  |  |  |
|  | 2 |  |  | 3.9 | 2.6 |
|  | 3 |  |  |  |  |
| 3'(2') 5'-bisphosphate nucleotidase CysQ, cysQ, Q1PWJ7 | 1 |  |  |  |  |
|  | 2 |  |  | 3.1 | 1.2 |
|  | 3 |  |  |  |  |

Relative content
2E-03
1E-03
5E-04
1E-04
1E-05
1E-06
Fold change

**Table 4 B:** Comparison of anammox proteins expressions in anammox biomasses adapted to low temperature (15°C) via cold shocks and via gradual acclimation. Anammox proteins upregulated more efficiently by cold shocks, excluding proteins already mentioned in Table 3. Sample 1 - day 0, sample 2 - distinct activities, sample 3 - end of experiment. Note: Red triangle in upper right corner…protein not detected, the value was set to detection limit 1E-6.

| Description, gene tag,<br>accession number | Sample | Relative<br>content |  | Fold change<br>compared to initial<br>state |  |
| --- | --- | --- | --- | --- | --- |
|  |  | Shocked | Gradual | Shocked | Gradual |
| Uncharacterized<br>protein, KSMBR1_1386,<br>Q1PZ66 | 1 |  |  |  |  |
|  | 2 |  |  | 2.8 | 2.5 |
|  | 3 |  |  |  |  |
| Uncharacterized<br>protein, KSMBR1_1565,<br>A0A2C9CE69 | 1 |  |  |  |  |
|  | 2 |  |  | 2.5 | 2.1 |
|  | 3 |  |  |  |  |
| Similar to<br>ribonucleoside<br>triphosphate reductase<br>NrdD, nrdD, Q1PZG5 | 1 |  |  |  |  |
|  | 2 |  |  | 2.3 | 0.2 |
|  | 3 |  |  |  |  |
| Strongly similar to<br>aspartate transaminase,<br>aspAT, Q1PXR8 | 1 |  |  |  |  |
|  | 2 |  |  | 1.7 | 1.0 |
|  | 3 |  |  |  |  |
| Uncharacterized<br>protein, KSMBR1_3326,<br>A0A2C9CJA4 | 1 |  |  |  |  |
|  | 2 |  |  | 1.5 | 1.4 |
|  | 3 |  |  |  |  |
| Uncharacterized<br>protein, kuste2346,<br>Q1Q690 | 1 |  |  |  |  |
|  | 2 |  |  | 1.5 | 1.4 |
|  | 3 |  |  |  |  |

To complement the proteins which changed significantly, we determined the expression of putative (already identified) cold shock proteins and other stress proteins, such as heat shock proteins and universal stress proteins. Table S 3 confirms that CspB played the dominant role in cold adaptation mechanism, as the other putative cold shock proteins (CspA - cold shock protein A, RecA - recombinase A, GyrA - gyrase A, GyrB - gyrase B, Sigma 54) were much less abundant and much less efficiently upregulated. The second most efficient putative cold shock protein was recombinase A (1.73-fold increase). Of the putative heat shock proteins, the cold shocks uniquely upregulated only a chaperone HtpG (high-temperature protein G) (1.6-fold increase), transmembrane isomerases ppiD and ppiC (Peptidyl-prolyl cis-trans isomerase D and C), and several small heat shock proteins.

Csp proteins in anammox have first been investigated in a study of several anammox populations exposed to temperature reduction from 35 to 25 °C; certain unspecified Csp proteins were most efficiently upregulated in “*Ca*. Brocadia fulgida” than in “*Ca*. B. sinica” and “*Ca*. Jettenia caeni”, potentially giving “*Ca*. B. fulgida” a metabolic advantage (Huo et al., 2020). This suggests that the upregulation of Csp proteins could be a promising mechanism of anammox adaptation to low temperatures.

Due to the lack of anammox-based metaproteomics studies, the following discussion of the function, structure and expression of Csp proteins is based on other bacteria. Our results suggest that CspB plays the most important role in the cold shock adaptation of “*Ca*. Kuenenia stuttgartiensis” (Table 3). CspB also played a major role in a study of *Bacillus subtilis*, in which the reduction of cultivation temperature from 37 to 10 °C resulted in a 20-fold upregulation of CspB, and mutant cells without CspB were less viable after freezing compared to parental cells (Willimsky et al., 1992). In comparison, our study showed the somewhat less efficient upregulation of CspB (5-to 12-fold increase), but this is similar to the 2-to 10-fold upregulation of dominant Csps reported for *Escherichia coli* (Goldstein et al., 1990). Structurally, CspB, like other Csp proteins, is highly conserved, relatively small (7.5 kDa), and consists of 5 antiparallel ß-strands forming a ß-barrel that adopts an oligonucleotide and/or oligosaccharide binding fold. Also like other major Csp proteins, CspB binds to DNA and RNA by the RNP1 and RNP2 nucleic acid binding motifs in order to destabilize undesirable secondary structures and maintain the nucleic acid structure in a single strand (Lindquist and Mertens, 2018). Under low temperatures and other sub-optimal conditions, single strands of nucleic acids form stable structures by base pairing, which is thought to inhibit transcription and translation. Csp proteins prevent this by functioning as nucleic acid chaperones, which are not to be confused with protein chaperones; protein chaperones not only destabilize undesirable secondary structures but also often restore the correct secondary and tertiary structures (Horn et al., 2007).

Our study also suggests that “*Ca*. Kuenenia stuttgartiensis” adapts to low temperatures by other proteins such as TypA (ribosome-binding GTPase), PpiD and UspA (Universal stress protein A) (Table 3, Table 4). TypA (also known as BipA) was found to possess chaperone activity and be essential for *E. coli* and *Pseudomonas putida* under low-temperature stress (Choi and Hwang, 2018; Pfennig and Flower, 2001; Reva et al., 2006). PpiD was also shown to function as a chaperone (Matern et al., 2010) and while its role has been studied under heat stress (Noor, 2015), our study provides the first evidence of its upregulation during cold stress (shocked 3.2-fold; gradual acclimation 1.4-fold). This dual role, acting as heat or cold shock proteins, in different organisms is well established; for example, DnaK (a typical heat shock protein / chaperone) was upregulated after cold shock in *Lactococcus piscium*. UspA in psychrotrophic *L. piscium* cultivated at 5 °C (growth optimum at 26 °C) increased 1.4-fold first with suspected cold shock proteins and subsequently 2.1-fold in the ‘cold acclimation’ phase (Garnier et al., 2010). This suggests that UspA, which is synthesized continuously during cultivation at low temperatures, acts as a cold acclimation protein whose upregulation starts later than Csp upregulation (Hébraud and Potier, 1999). In comparison, our study recorded the 2.6- and 1.5-fold upregulation of UspA in shocked and gradually acclimated reactors, respectively, suggesting the more efficient adaptation of shocked biomass to low temperature.

It is reasonable to assume that these Csps re-started protein synthesis, including the synthesis of N-respiration proteins (e.g. NXR, HZS,…) responsible for anammox activity; thus, we suggest that more efficient induction of Csps by cold shocks compared to gradual acclimation made anammox cells more active.

### 3.3 Structural changes in ladderane lipids of “*Ca*. Kuenenia” under reduced temperature

UPLC-MS/MS revealed that cold shocks and gradual acclimation induced changes in ladderane composition (Table 5). The alkyl moieties were altered in two main ways. First, a significant decrease in the length of the [5]-ladderanes shifted the C20/(C18+C20) ratio in favor of C18 (control 0.49; shocked 0.33; acclimated 0.31). Second, of the two ladderane esters and/or ethers on the glycerol backbone, the one in the sn-1 position was replaced by a straight or branched non-ladderane alkyl (C14/C15/C16: control 35%; shocked 55%; acclimation 51%). These replacements were exclusively C14 or C16, and the resulting lipids comprised 25-28% (sn positions: 1 - C14 ester; 2 - C20-[5]-ladderane) and 17-20% (1 - C16 ester; 2 - C20-[5]-ladderane) of all ladderane lipids, respectively. Concerning the polar headgroup, the content of phosphatidylcholine (PC) and phosphatidylglycerol (PG) was reduced in favor of phosphatidylethanolamine (PE) (PE: control 15%; shocked 25%; acclimated 24%).

**Table 5:**
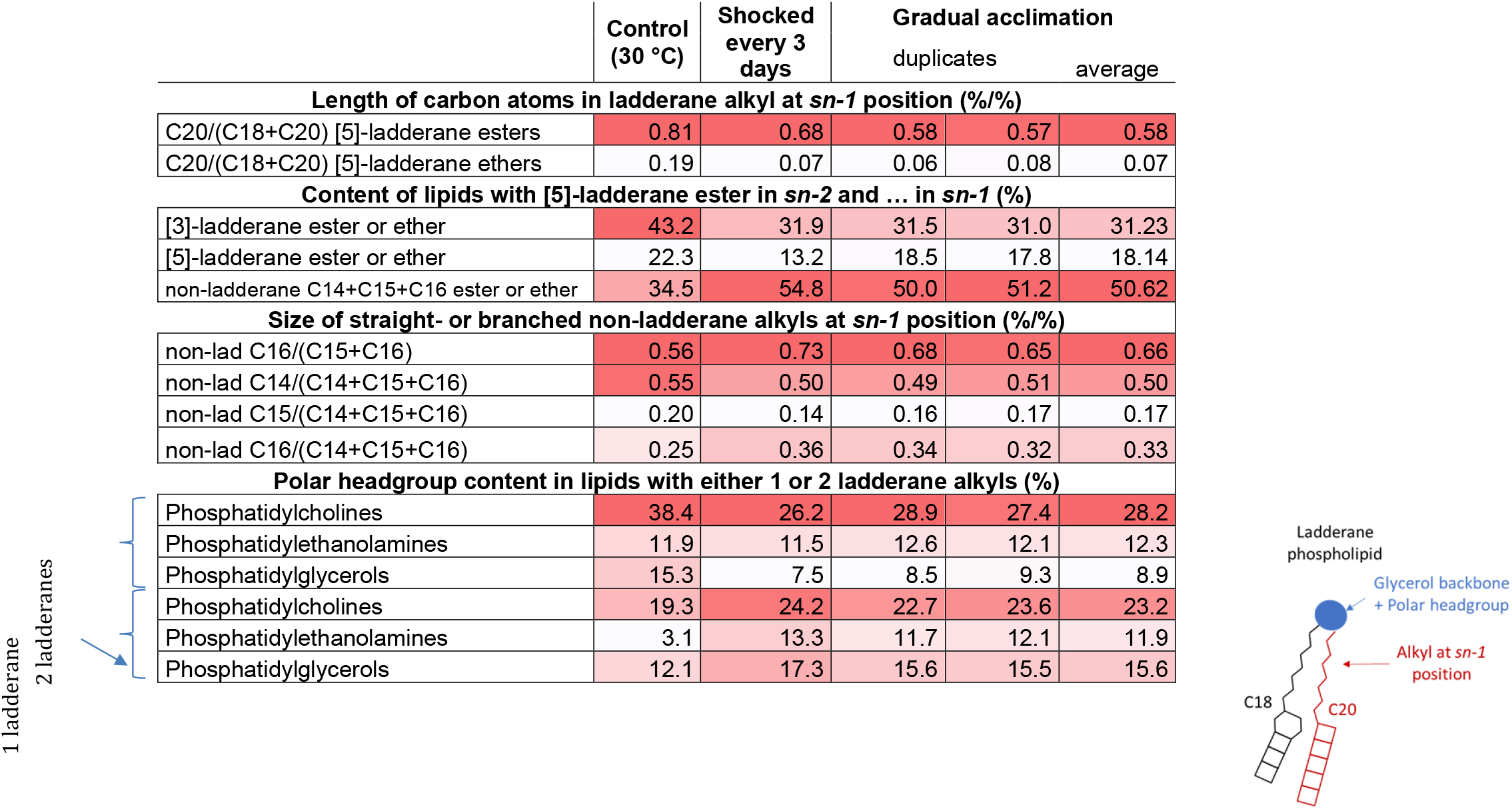
Relative content (based on lipid signal cps) of ladderane lipids in anammox biomass exposed to cold shocks (day 44), gradual acclimation (day 46) and the control culture kept at 30 °C. Ladderane lipid content in anammox biomasses revealed that cold shocks and gradual acclimation induced the synthesis of shorter [5]-ladderanes, replacement of [5]-ladderanes on *sn-1* position by C14 and C16 non-ladderane alkyls and phosphatidylethanolamine as polar headgroup. The color gradient highlights more prevalent membrane lipid components. Note: [5]-ladderanes…5 concatenated cyclobutane rings; C20, C18…number of C atoms in the respective alkyl

To sum up, while the adopted sampling regime does not enable us to attribute changes in ladderane composition to increased activity of the shocked culture, we can state that “*Ca*. Kuenenia” adapts to low temperatures by synthesizing shorter ladderanes, by replacing ladderanes with even shorter alkyl esters or ethers, and by favoring PE as the polar headgroup. Thus, our results match with the fact that bacteria prevent membrane rigidity at lower temperatures by modulating their membrane lipidic composition and fluidity via mechanisms such as synthesizing shorter and more branched alkyls (Siliakus et al., 2017). The increased prevalence of PE as the relatively smaller polar headgroup may ease the insertion of membrane proteins that could enable further cold adaptation (van Klompenburg et al., 1998).

### 3.4 Strategies for adaptation of anammox bacteria to low temperatures

Compared to the unadapted “*Ca*. Kuenenia” (Kouba et al., 2019), cold shocks (5 °C, 1 h) and gradual adaptation (1 °C/day) increased the anammox activity by 130% and 40%, respectively (shocked once per 3 days 0.74±0.15 kg-N/kg-VSS/d; gradually acclimated 0.45±0.06 kg-N/kg-VSS/d; unadapted culture 0.32±0.019 kg-N/kg-VSS/d; 15 °C), suggesting cold shocks to be the superior strategy for low temperature adaptation. In the literature, gradual adaptation (De Cocker et al., 2018) along with the enrichment of cold-adapted species (Hendrickx et al., 2014) are the established strategies for achieving high-performance anammox at low temperatures (10-20 °C), while the cold shocks have only very recently emerged as a promising alternative. Our previous work in batch assays has shown that the application of 5 °C for 8 h consecutively elevated the anammox activity at 10 °C (Kouba et al., 2018), while we have subsequently demonstrated that the elevated activity of one such shock can endure for 40 days (Kouba et al., accepted). In our present work, we reported a shorter duration (14-18 days) of increased activity post-shock. The shorter duration in our study may be explained by a shorter exposure to 5 °C (present study 1 h; Kouba et al. accepted 8 h).

On a practical note, increasing the process resiliency against low temperature would require the application of the present cold shock regime once every 6 weeks, which means that a 6-month winter period would require the application of 4-5 such regimes. The shock applications could be made less frequent by optimizing cold shock parameters such as shock duration and temperature which have the potential to extend the time-span of the increased resiliency post-shock. The anammox sludge could also be shocked in the late summer and autumn to prepare the bacteria for the winter season in a side reactor cooling down a concentrated anammox sludge (e.g. return sludge line). It can also be applied to the mesophilic side-stream cultures before the inoculation of main-stream facilities (Kouba et al., accepted). Overall, this is an important step in establishing cold shocks as a viable adaptation strategy.

## 4 Conclusions

Mainstream anammox operations for an energy- and resource-efficient removal of nitrogen from municipal wastewater requires an in-depth understanding of mechanisms of microbial adaptations to temperatures across seasons, notably in the perspective of process control ahead of winter conditions. We showed that:

- Anammox enrichment cultures of “*Ca*. Kuenenia stuttgartiensis” exposed to cold shocks (5 °C, 1 h shock duration, shocked once every 3 or 7 days) adapted to subsequent operation at low temperature (15 °C) more favorably (0.66±0.06 kg-N/kg-VSS/d) compared to a gradually acclimated culture (−1 °C/day) (0.48±0.02 kg-N/kg-VSS/d) for the duration of 14-18 days post-shock.
- The increased activity of shocked cultures was linked to (i) maintaining the levels of proteins involved in nitrogen respiration and (ii) upregulation of several putative cold shocks proteins (e.g. CspB, TypA, PpiD) and of several hypothetical ones that should be better characterized.
- Low-temperature adaptation resulted in several significant changes in the structure of characteristic membrane ladderane lipids, such as higher content of shorter (C18) ladderanes and even shorter (C14-16) straight/branched alkyls, and PE as the polar headgroup.

Overall, the results underline the efficiency of cold shocks as a promising anammox adaptation strategy for mainstream operations for nitrogen removal from municipal WWTP. We identified the yet unreported physiological mechanisms of adaptation of anammox microorganisms under cold shocks and gradual acclimation, that underlie an increased anammox activity at low temperature.

## 5 Acknowledgements

The authors acknowledge the financial support of the Czech Ministry of Education Youth and Sports through project GACR 17-25781S. Dana Vejmelkova was supported by a Talent grant of the Soehngen Institute of Anaerobic Microbiology (SIAM, project 62002148) which is financed by a Gravitation grant from the Dutch Ministry of Education, Culture and Science. Internal partial funding from the TU Delft is acknowledged. Michele Laureni was supported by a Marie Skłodowska-Curie Individual Fellowship (grant agreement 752992) and a VENI grant from the Dutch Research Council (NWO) (project number VI.Veni.192.252).

## 7 Supplementary materials

The picture of experimental set-up is depicted at Fig. S 1.

**Fig. S 1:**
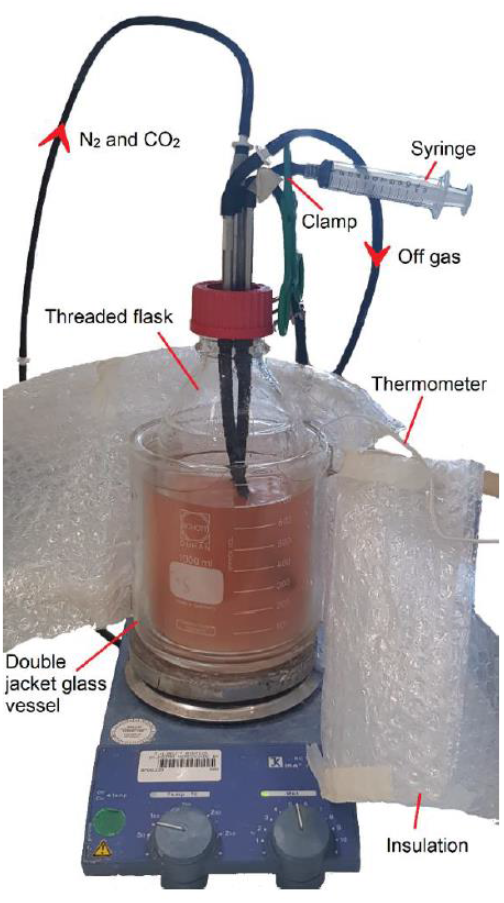
experimental set-up.

The composition of the cultivation medium fed to the anammox reactors is described in Table S 1 and 2.

**Table S 1:** Composition of feed to fed batch reactor for planktonic “*Ca*. Kuenenia”. Fe2+ solution was prepared by dissolving 9.14 g of FeSO4.7H2O and 6.37 g EDTA in 1 L of Milli-Q water, with pH 2.5. Mg2+ solution was prepared by dissolving 160 g of MgSO4.7H2O in 1 L of Milli-Q water. Ca2+ solution was prepared by dissolving 240 g CaCl2.2H2O in 1 L of MilliQ water. Preparation of trace elements solution is described in Table S 2. The pH of final solution is adjusted to 7.1 to 7.2 by adding 4M NaOH or 1M H2SO4. Nitrite and ammonium were added in separate solutions.

| Substance / solution | Units | Value |
| --- | --- | --- |
| K<sub>2</sub>HPO<sub>4</sub> | [g] | 18.231 |
| KH<sub>2</sub>PO<sub>4</sub> | [g] | 2.60 |
| Trace Element | [mL] | 31.25 |
| Fe<sup>2+</sup> | [mL] | 62.50 |
| Mg<sup>2+</sup> | [mL] | 15.625 |
| Ca<sup>2+</sup> | [mL] | 7.8125 |

**Table S 2:** Composition of trace elements solution for the preparation of feed for anammox reactors per 1 L of Milli-Q water and pH of the final solution is adjusted to 6 using solid NaOH.

| Substance / solution | Units | Value |
| --- | --- | --- |
| EDTA | [g] | 19.11 |
| ZnSO<sub>4</sub>·7H<sub>2</sub>O | [g] | 0.43 |
| CoCl<sub>2</sub>·6H<sub>2</sub>O | [g] | 0.24 |
| MnCl<sub>2</sub>·4H<sub>2</sub>O | [g] | 1.0 |
| CuSO<sub>4</sub>·5H<sub>2</sub>O | [g] | 0.25 |
| (NH<sub>4</sub>)<sub>6</sub>Mo<sub>7</sub>O<sub>24</sub>·4H<sub>2</sub>O (=1.25 mM Mo) | [g] | 0.22 |
| NiCl<sub>2</sub>·6H<sub>2</sub>O | [g] | 0.20 |
| HNaSeO<sub>3</sub> | [g] | 0.09 |
| H<sub>3</sub>BO<sub>3</sub> | [g] | 0.014 |
| Na<sub>2</sub>WO<sub>4</sub>·2H<sub>2</sub>O | [g] | 0.054 |

### Reagents for protein extraction

TEAB (triehtylammonium bicarbonate resuspension buffer) - 50 mM TEAB, 1% (w/w) NaDOC, pH = 8.0 by HCl

NaDOC (Sodium Deoxycholate, Sigma-Aldrich)

B-PER buffer

100% (w/v) Trichloroacetic acid (TCA), dissolve 500g TCA in 350 mL dH2O, store at RT.

**Table S 3:**
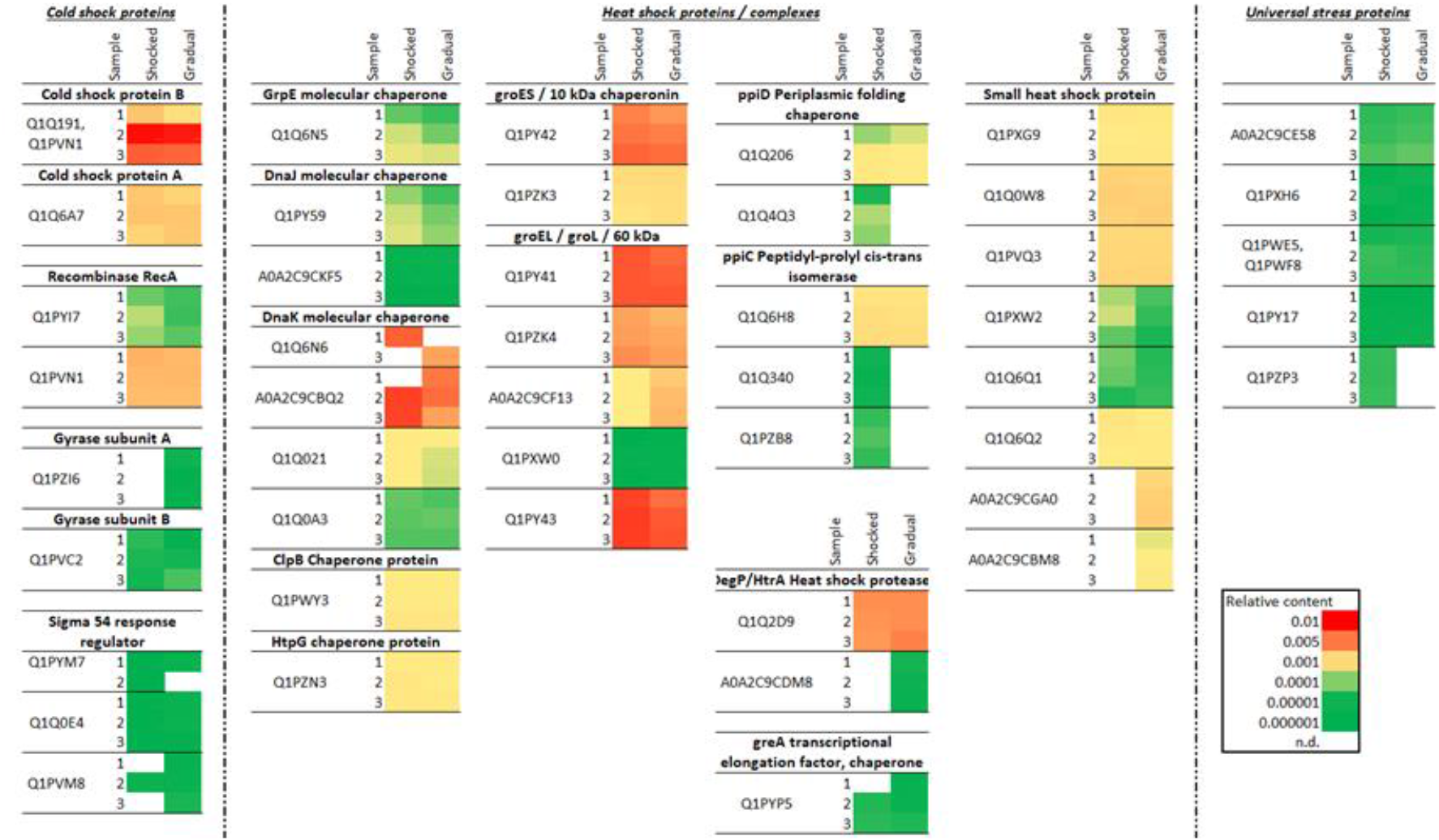
Relative content of putative cold and heat shock proteins in anammox biomasses exposed to cold shocks and to gradual acclimation to 15°C. Most pronounced upregulation was observed for CspB. PpiC, PpiD and parts of chaperon complex GrpE-DnaJ-DnaK-ClpB-HptG were more efficiently upregulated by cold shocks compared to gradual acclimation. Sample 1 - day 0, sample 2 - distinct activities, sample 3 - end of experiment.

## Notes

### Competing Interest Statement

The authors have declared no competing interest.

